# Quantifying relative sponginess: a high-resolution model of landscape water retention as an ecosystem service

**DOI:** 10.1101/2024.10.11.617771

**Authors:** Paul M. Evans, Alejandro Dussaillant, Varun Varma, John W. Redhead, Richard F. Pywell, Jonathan Storkey, Andrew Mead, James M. Bullock

## Abstract

Unprecedented climate and land use changes are having major impacts on water-based ecosystem services (ES). It is crucial, therefore, to get an in-depth understanding of current levels of such ES provision, and how they could be impacted by changed environments or management. However, applying existing models for water-related ES pose substantial challenges, which include the need for in-depth specialist hydrological knowledge, the requirement for numerous datasets and parameters that may not be consistently available, and high computational costs. Additionally, there is often a mismatch between the resolution of the model output and parcel-level land management, at which ES information is often most valuable for supporting decision-making.

Here we detail a rapid method for assessing water retention potential by estimating an area’s ‘sponginess’ - its capacity to absorb and retain precipitation. Our approach builds upon a topography-adjusted Curve Number methodology, a widely recognised and straightforward tool for estimating water run-off. We calculated run-off across 1 km grid cells (representing field- to farm-scale land parcels) in Great Britain based on the land’s sponginess under storm conditions. The primary objective was to investigate the applicability of the method in estimating point-values — i.e., the independent contribution of each grid square — within the context of known limitations. The results enable the identification of areas with higher potential for the ES of water retention.

Our model output illustrates the spectrum of sponginess across Great Britain, ranging from less than 30% of precipitation retained in city regions to as high as 99% in some rural, agricultural areas. Importantly, we demonstrate that the model is easy to run, can be used with freely-available data, and produces outputs compatible with grid-based models for other ES. Overall, the model provides an accessible approach to estimating the ES of water retention to researchers worldwide, even in data-scarce areas.

**Highlights:**

- To inform land management decisions it is important to determine parcel-level ecosystem services, but there is a gap for services related to water.
- Parcel-level assessments of a land’s ‘sponginess’ – the capacity to retain precipitation – are described.
- Values are calculated using a topography-adjusted Curve Number methodology, which requires little data input or hydrological expertise.
- ‘Sponginess’ and run-off estimates are provided for Great Britain at 1 km resolution.

## 1. Introduction

Management of water resources is becoming increasingly important due to a changing climate and urbanisation creating more impervious land, with both droughts and floods forecast to increase in frequency and intensity globally (Alfieri et al., 2015; Spinoni et al., 2015). Severe storm events are also more likely (Cotterill et al., 2021). Recent floods alone have had a major impact, comprising almost half of weather-related disasters, and affecting around 2.3 billion people over the 10 years to 2015 (UNDRR, 2015). Flooding can be rapid, and can have major health, infrastructure, environmental and biodiversity impacts (e.g., Talbot et al., 2018; Suhr and Steinert, 2022). Reducing the probability of flooding is therefore highly beneficial, and is considered a regulating Ecosystem Service (ES), often defined as ‘flood control’, ‘flood regulation’ or ‘water retention’ (Vári et al., 2022). Given that the importance of this ES will only increase because of climate change, it is imperative that its response to land use change (which may be driven by other priorities such as house-building targets) is adequately captured in planning decisions.

However, the mapping of any ES relating to water flow can be extremely challenging, often demanding extensive, fine-scale input datasets and the parameterisation of complex numerical models (e.g., Bell et al., 2009; Douglas-Mankin et al., 2010; Kay et al., 2023). Such models achieve high precision but also come with a very high resource cost in terms of computational power, time requirements, and expertise, which may be prohibitive when making land management decisions at the catchment or landscape scale. Therefore, approaches that require fewer resources to run and are simpler to use, but importantly, are well founded in hydrological science, are required to address this issue. These requirements are especially important for data-scarce regions (Benra et al., 2021), and for performing quick assessments based on land change scenarios. Simpler ES models have been produced to address such constraints (e.g., Borota et al., 2024). Sharps et al. (2017) compared three such simple but effective ES models for water supply. One model mentioned in Sharp et al. (2017) has been used over 600 times across the world including in regions where the capacity to run more complex models is low. In such regions, training with simple models can be extremely advantageous (Bhagabati et al., 2014; Ruckelshaus et al., 2015; Benra et al., 2021). Simple models provide spatially uniform estimates, have a user-defined resolution, and are relatively quick to run.

Furthermore, models that produce uniform per-grid square values have the advantage that they can readily be used in assessments of trade-offs and synergies with other ES at individual locations, where these ES are hydrological (Duku et al., 2015) or non-hydrological (Remme et al., 2014; Aryal et al., 2022). This ability to compare ES is important as, for example, policy targets set by institutions such as the UK and European Parliaments aim to enhance multiple ES and thus transition towards multifunctional landscapes (van Zanten et al., 2014; POST, 2021). In such multifunctional landscapes, water retention may not be the main aim, but will be affected by decisions made (e.g., the UK’s Sustainable Farming Incentive, whose main aims align with supporting food production (UK Government, 2024)). Such trade-offs can be difficult to determine from more complex water ES models, as they often focus on aggregated measures of water processes at the catchment scale and depend on the influence of upstream areas. Therefore, models that more closely match the specific details (i.e., resolution and extent) of individual land parcels are useful when estimates are needed at the spatial scale at which land is managed. This can aid tailored approaches to landscape management across varying scales (Stürck and Verburg, 2017), and for divergent interests of decision makers (Hölting et al., 2020; Sun et al., 2020; Aryal et al., 2022).

Each land parcel, whether an extensive arable field or a small garden, possesses an inherent capacity to retain water, whether this water comes in the form of precipitation or run-off from neighbouring land parcels. This property, termed “sponginess,” influences the amount of precipitation that runs off. Sponginess can be thought of as a function of the land parcel’s land use composition, among other factors, and can, therefore, be affected by land management practices (e.g., Erena and Worku, 2019; Srivastava et al. 2020; Abdallah et al., 2021). The lower run-off under higher sponginess is due to the greater amount of water exiting the area only through evaporation and evapotranspiration, and the capacity to hold water, leading to key hydrological functions such as infiltration, percolation and replenishment of groundwater. Sponginess can be divided into contributions from “green,” “blue,” and “grey” infrastructure, referring to green, vegetated spaces; water bodies; and built infrastructure designed to help deal with stormwater, respectively (e.g., Zhang et al., 2019; Zuniga-Teran et al., 2020; ARUP, 2023).

In this study, the sponginess of all 1 km grid squares in Great Britain was calculated using the Curve Number (CN) method (Mockus, 1949; USDA, 2004) to capture existing gradients at the national scale and facilitate benchmarking within prescribed landscapes. The CN method is a widely used, simple, empirically-based, and effective tool for estimating run-off in watershed modelling, hydrological analysis, and water resources management (Jones, 1997; Shaw et al., 2011). Initially created to estimate surface run-off depth in small catchments given a specified rainfall event (Mockus, 1949; USDA, 2004), it has since been expanded and improved to encompass non-agricultural watersheds and used in a wide range of applications (e.g., Hawkins et al., 2019; Młyński et al., 2020; Soulis, 2020, and references therein). Latterly, the inclusion of slope, which was omitted from earlier calculations, has been shown to be very important (e.g., Sharma et al., 2022). Therefore, in this study, a topographic slope-adjusted CN method was employed, for the first time to our knowledge, to estimate the relative water retention of grid squares following storm events for a whole country, thus estimating the land’s sponginess. Our primary objective was to investigate the applicability of the method, and its utility for identifying areas with a higher potential for water retention and reduced run-off potential, particularly within the context of constraints on resources, computational ability, time requirements, and expertise.

## 2. Methods

### 2.1. Curve Number method

The Curve Number (CN) method is briefly explained below, and in more detail elsewhere (e.g., Lian et al., 2020). CN uses the water balance equation of a closed system, which states that any water that flows into an area is equal to outflow plus storage in fractured rocks, soils, and groundwater (Mishra and Singh, 1999), represented in the general CN run-off equation (eq. 1).

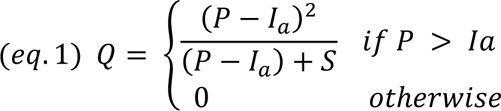

where *Q* is the depth of run-off i.e., the amount of water that leaves a grid square downstream, *P* is the depth of rainfall received, *I_a_* is the initial abstraction, mainly from available canopy interception, surface depression storage and initial infiltration, and *S* is the maximum potential retention. All units are mm.

For this model, the only water input to a grid square is precipitation, and the outflows can include evaporation, evapotranspiration, and discharge (Jones, 1997; Shaw et al., 2011). Thus, each area does not receive any run-off from upslope processes. This allows comparative metric of an area’s ability to retain water - its sponginess – to be derived. The main parameter required is the CN itself, which is derived from a combination of hydrologic soil group (HSG), land cover, and the Antecedent Moisture Condition (AMC); the last refers to the moisture condition at the beginning of a rainfall event. For this study, a moderate AMC was used as that is standard practice; however, other options are available (see Supplementary Information, S1). The CN is a transformation of *S,* the maximum potential retention (eq. 2a, 2b), which makes interpolation more linear (USDA, 2004).

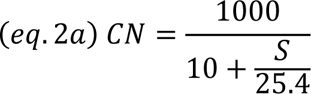

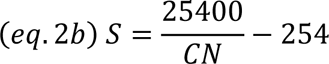

where *CN* is the Curve Number, a dimensionless parameter with the range from 0 to 100 (where low CN means more storage thus less runoff); and *S* is the maximum potential retention (mm).

### 2.2. Curve Number values

The original CN values were derived in 1986 from data collected in the USA, the tables of which can be found in USDA (2004), commonly referred to as TR-55. While there have been some studies that have used different values, most research still focuses on the values from TR-55. One study that used different values for the UK (specifically for Pickering in North Yorkshire) was Thomas and Nisbet (2016).

Values for this study were derived mainly from TR-55, and supplemented with Thomas and Nisbet (2016) values where the former study did not include specific land cover types (Table 1). TR-55 proposes different CN values based on the land cover condition and hydrologic soil group (HSG). Following methodology in Hooftman et al. (2023), a 2015 25 x 25 m resolution land cover map (Rowland et al., 2017) was combined with Land Cover Plus Crops for 2016 (https://www.ceh.ac.uk/data/ceh-land-cover-plus-crops-2015), producing 31 land cover types that were used in this study. All land cover types were assumed to be in ‘good’ condition, except for acid grassland which was considered as ‘Bracken’ (i.e., ‘fair’ condition) from Thomas and Nisbet (2016). The HSG data used was 250 x 250 m spatial resolution (Ross et al., 2018). Land cover and HSG were overlaid, and a CN obtained for each 25 x 25 m cell.

**Table 1:** Curve number (CN) for land cover and hydrologic soil group (HSG) combinations for the UK.

| Land cover (condition) | Land cover classification | Treatment | Curve numbers for hydrologic soil group |  |  |  |
| --- | --- | --- | --- | --- | --- | --- |
|  |  |  | A | B | C | D |
| Woods (good)<sup>1</sup> | Broadleaf woodland | N/A | 30 | 55 | 70 | 77 |
| Woods (good)<sup>1</sup> | Coniferous woodland | N/A | 30 | 55 | 70 | 77 |
| Rotation meadow (good)<sup>2</sup> | Arable - grass | straight row | 67 | 78 | 85 | 89 |
| Small grain (good)<sup>2</sup> | Arable – winter wheat | straight row | 63 | 75 | 83 | 87 |
| Small grain (good)<sup>2</sup> | Arable – winter barley | straight row | 63 | 75 | 83 | 87 |
| Small grain (good)<sup>2</sup> | Arable – spring wheat | straight row | 63 | 75 | 83 | 87 |
| Small grain (good)<sup>2</sup> | Arable - oats | straight row | 63 | 75 | 83 | 87 |
| Row crops (good)<sup>2</sup> | Arable - maize | straight row | 67 | 78 | 85 | 89 |
| Row crops (good)<sup>2</sup> | Arable – oil seed rape | straight row | 67 | 78 | 85 | 89 |
| Small grain (good)<sup>2</sup> | Arable – spring barley | straight row | 63 | 75 | 83 | 87 |
| Row crops (good)<sup>2</sup> | Arable - potato | straight row | 67 | 78 | 85 | 89 |
| Broadcast legumes (good)<sup>2</sup> | Arable - beans | straight row | 58 | 72 | 81 | 85 |
| Row crops (good)<sup>2</sup> | Arable - other | straight row | 67 | 78 | 85 | 89 |
| Pasture, grassland (good)<sup>1</sup> | Improved grassland | pasture | 39 | 61 | 74 | 80 |
| Meadow (-)<sup>3</sup> | Neutral grassland | N/A | 30 | 58 | 71 | 78 |
| Meadow (-)<sup>3</sup> | Calcareous grassland | N/A | 30 | 58 | 71 | 78 |
| Brush (fair)<sup>4</sup> | Acid grassland | N/A | 35 | 56 | 70 | 77 |
| Meadow (-)<sup>3</sup> | Fen, marsh and swamp | N/A | 30 | 58 | 71 | 78 |
| Brush (good)<sup>5</sup> | Heather | N/A | 30 | 48 | 65 | 73 |
| Meadow (-)<sup>3</sup> | Heather grassland | N/A | 30 | 58 | 71 | 78 |
| Meadow (-)<sup>3</sup> | Bog | N/A | 30 | 58 | 71 | 78 |
| Open water (-)<sup>6</sup> | Inland rock | N/A | 100 | 100 | 100 | 100 |
| Open water (-)<sup>6</sup> | Saltwater | N/A | 100 | 100 | 100 | 100 |
| Open water (-)<sup>6</sup> | Freshwater | N/A | 100 | 100 | 100 | 100 |
| Bare soil (-)<sup>7</sup> | Supralittoral rock | N/A | 77 | 86 | 91 | 94 |
| Bare soil (-)<sup>7</sup> | Supralittoral sediment | N/A | 77 | 86 | 91 | 94 |
| Bare soil (-)<sup>7</sup> | Littoral rock | N/A | 77 | 86 | 91 | 94 |
| Bare soil (-)<sup>7</sup> | Littoral sediment | N/A | 77 | 86 | 91 | 94 |
| Open water (-)<sup>6</sup> | Saltmarsh | N/A | 100 | 100 | 100 | 100 |
| Impervious areas<sup>8</sup> | Urban | N/A | 89 | 92 | 94 | 95 |
| Fair condition urban (grass cover 50% to 75%)<sup>9</sup> | Suburban | N/A | 49 | 69 | 79 | 84 |

Note: 'Ref 1' and 'Ref 2' indicate that values were obtained from Thomas and Nisbet (2016) and TR-55 (USDA, 2004), respectively. <sup>1</sup>Ref 2 used because it had 'good' values; <sup>2</sup>Ref 2 used due to assumption the fields would not be fallow; <sup>3</sup>Set-a-side grassland' value used from Ref 1 (i.e., assumed to be free from grazing); <sup>4</sup>'Bracken' value used from Ref 1; <sup>5</sup>'Brush (good)' value used from Ref 2; <sup>6</sup>'Open water' value used from Ref 1; <sup>7</sup>'Bare ground' value used from Ref 1; <sup>8</sup>'Urban' value used from Ref 1; <sup>9</sup> Ref 2 values used because it was lower.

### 2.3. Slope

The original CN method USDA (2004) did not account for slope because the equations were derived from American farms with gradients of less than 5%. Consequently, the slope of various research locations was not considered a significant factor, and the originally tabulated CN values were applied as standard practice regardless of gradient (Lal et al., 2017). However, slope can have a substantial effect on run-off, and it is therefore important to include this parameter in calculations for regions with greater topographic variation (Sharpley and Williams, 1990; Sharma et al., 2022). Moreover, absolute slope per cent is more accurate than categorised versions (e.g., Sharpley and Williams 1990; Huang et al., 2006; Ajmal et al., 2016). Here, we used the Sharpley and Williams (1990) equation for slope-adjusted CN (eq. 3) as Sharma et al. (2022) determined that was the most applicable for general CN calculations where slope was included.

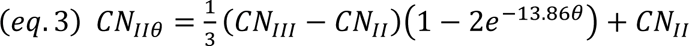

where *CN*_IIθ_ is the slope-adjusted CN for normal conditions (i.e., moderate AMC), *CN_II_* is the tabular CN value for normal conditions, *CN_III_* is the tabular CN value for wet conditions, and *θ =* average land slope (m/m) for a grid square (originally ‘average land slope (m/m) of the catchment’ (Sharpley and Williams 1990)). Here, slope was calculated using 50-metre Integrated Hydrological Digital Terrain Model output (Morris et al., 1990).

### 2.4. Initial abstraction (I_a_) values

The initial abstraction (*I_a_*) is an important parameter that accounts for precipitation losses prior to run-off. Such losses consist mainly of canopy interception - precipitation collected on leaves, stems, and branches and evaporated back into the atmosphere before reaching the ground (Kermavnar and Vilhar, 2017), early infiltration into the soil depending on soil condition, and the maximum surface depression storage e.g., roughness ruts or pits (NRCS, 2004). In its general form, *I_a_* is a variable fraction of *S,* determined using an initial abstraction ratio coefficient, λ (USDA, 1986; eq. 4).

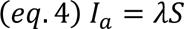

The default λ value is 0.20 (USDA, 1986); however, various estimates ranging from 0 to 0.86 have been suggested to improve accuracy, depending largely on the region of study (see Ling and Yusop (2014) and references therein). Ideal λ values were not known for GB, therefore we approximated an initial abstraction ratio measure for each land use type guided by two important *I_a_* parameters, canopy interception and a proxy for surface depressions. The latter was determined using definitions from Acrement and Schneider (1984) about how different land use including depressions affects water flow (herein ASV) (see Supplementary Information, S2).

First, we gave the GB land covers with the highest known rainfall interception, woodlands (Broadmeadow et al., 2023), the highest λ values. In the UK, research shows that annual woodland canopy interception is between 25% – 45% and 10% – 25% of total rainfall for coniferous and broadleaved woodlands, respectively (Nisbet et al., 2005). Using modelled counterfactual scenarios, Broadmeadow et al. (2023) determined that the mean daily interception loss for ‘storm’ days (classed as > 25 mm rainfall) was 6.47 and 3.05 mm for coniferous and broadleaved woodlands, respectively. Using these values, it can be inferred, assuming a 25 mm storm, that coniferous woodland intercepted between 11% - 53%, with a mean of 26% of rainfall. For broadleaved woodland, the values were between 4% - 28% for a 25 mm rainfall event, with a mean of 12%. When combining that information with the values from Nisbet (2005) (i.e., from Fig. 2 in Nisbet (2005)), similar trends were identified. These mean values were used as the λ for woodland land cover types (i.e., 0.26 and 0.12 respectively for coniferous and broadleaved woodlands).

There were a paucity of data regarding non-tree canopy interception; however, leaf area index (LAI) is positively linked with canopy interception, especially for non-tree vegetation (Zheng et al., 2018; Nazari et al., 2020; Wang and Guo, 2024). Therefore, LAI – a proxy measure of canopy interception – was used in combination with ASV values to create relative vegetation λ indices (see Supplementary Information, S3).

LAI changes across the course of a year as plants grow. Daily storms (> 25 mm) occur more often in the winter in the UK (Tanguy et al., 2021), therefore we modelled the abstraction ratio coefficient as a function of winter LAI. Summer LAIs for European coniferous and broadleaf woodland were 4.65 and 4.74 m^2^m^−2^, respectively (Munier et al., 2018). In winter, the LAI of broadleaved (i.e., deciduous) trees can be greatly reduced (e.g., Potithep et al., 2013), although the woody stem and branches still cause interception, as does the leafy undergrowth. For these reasons, we halved the LAI to an approximate winter ‘LAI’ of 2.37 m^2^m^−2^ for broadleaf woodland. When comparing the winter LAI and the ASV index values for coniferous and broadleaf woodland, the latter was approximately half of the former (ASV = 0.2 and 0.1, and LAI = 4.65 and 2.37), meaning they were proportionally similar values (Table 2). Using the relationship between LAI and initial abstraction ratio coefficient for woodlands, a linear equation was derived (Eq. 5).

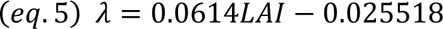

**Table 2:** Interception proportion values for 31 land cover types and the relative comparison values for proxy winter LAI values, and Acrement and Schneider’s (1984) values.

| Land cover | Rainfall interception proportion value ($\lambda$) | Assigned winter LAI ($\text{m}^2\text{m}^{-2}$) | Acrement and Schneider value (ASV) |
| --- | --- | --- | --- |
| Broadleaf woodland | 0.12 | 2.37 | 0.1 |
| Coniferous woodland | 0.26 | 4.65 | 0.2 |
| Winter wheat [winter sown] | 0.14 | 2.63 | 0.025 - 0.1 |
| Winter barley [winter sown] | 0.08 | 1.79 |  |
| Oil seed rape [winter sown] | 0.13 | 2.53 |  |
| Spring wheat [spring sown] | 0.04 | 1.09 |  |
| Oat [spring sown] | 0.05 | 1.25 |  |
| Maize [spring sown] | 0.03 | 0.85 |  |
| Spring barley [spring sown] | 0.03 | 0.92 |  |
| Potato [spring sown] | 0.06 | 1.35 |  |
| Beans [spring sown] | 0.10 | 2.10 |  |
| Other arable | 0.07 | 1.60 |  |
| Arable grass | 0.05 | 1.27 | 0.01 |
| Improved grassland | 0.05 | 1.27 |  |
| Neutral grassland | 0.12 | 2.44 |  |
| Calcareous grassland | 0.12 | 2.44 |  |
| Acid grassland | 0.12 | 2.44 |  |
| Heather grassland | 0.12 | 2.44 |  |
| Heather | 0.12 | 2.44 | 0.011-0.025 |
| Fen, marsh and swamp | 0.95 | N/A |  |
| Bog | 0.95 |  |  |
| Inland rock | 0.08 |  |  |
| Saltwater | 0.95 |  |  |
| Freshwater | 0.95 |  |  |
| Supralittoral rock | 0.08 |  |  |
| Supralittoral sediment | 0.4 |  |  |
| Littoral rock | 0.08 |  |  |
| Littoral sediment | 0.4 |  |  |
| Saltmarsh | 0.95 |  |  |
| Urban | 0.01 |  |  |
| Suburban | 0.03 |  |  |

The crop land cover types included in this study were identified as either winter or spring sown. Most LAI literature focuses on plants when they are in ‘full’ leaf, therefore, assuming storms mostly happen in winter, approximate LAI were assigned based proportional reductions from recorded full-leaf LAI (see Table A5.3 for LAI references). For winter-sown crops, which are generally sown in September or October, LAI was reduced by a quarter, as the crops are not fully in-leaf in winter. For spring-sown crops, which can be sown from March to May, LAI was reduced by a half to reflect that these crops are less developed (or even absent) throughout winter (Boardman et al., 2009; Vogel et al., 2016). Maize and potato were further reduced, in total to 25% of LAI, due to these crops having a high risk of run-off due to wide spacing and compaction when harvesting (Evans, 2005; Boardman, 2013; Environment Agency, 2024). This resulted in the crops having an average approximate LAI of 2.31 m^2^m^−2^, and 1.31 m^2^m^−2^ for winter and spring crops, respectively, equalling 12% and 7% interception (i.e., λ of 0.12 and 0.07) (Table 2). This is also reflected in the ASV, where cropland was equivalent to 12.5% to 50% of the maximum ASV. Equation 5 was also applied to arable, improved, and semi-natural grasses land cover types. For arable grasslands (i.e., leys) and improved grasslands, a winter effect was applied, halving the recorded LAI to an approximate winter LAI of 1.27 m^2^m^−2^ (5% interception), which has previously been recorded in the winter (e.g., Dusseux et al., 2014). LAIs for semi-natural grasslands were reduced by a quarter, resulting in 2.44 m^2^m^−2^ (12% interception) due to fewer flowers and leaves in the winter, but still a high architectural diversity for canopy interception. Data were lacking for LAI for heathland, so were assigned the same LAI as semi-natural grasslands.

Non-vegetated land cover λ values were based largely on associated surface depression storage. Rocky land covers, which can have deep depressions that are able to store water, were assigned a relatively modest 8% interception (i.e., λ of 0.08), although they can be much higher (Hu et al., 2020). Due to high porosity, sedimentary land covers, such as sand, were assigned 40% (Curry et al., 2004). Urban land cover was assigned a relatively low value of 1% as it is dominated by impervious surfaces which also hold very little water. Suburban was assigned 3% due to the slightly increased cover of gardens and green spaces.

### 2.5. Storm simulation

Here, we used the definition from Broadmeadow et al. (2023) of a storm in the UK, which is at least 25 mm of precipitation in a single event. To simulate different intensities of storm events three precipitation depths were used: 25 mm, 50 mm, and 75 mm. The primary focus was on the lowest storm simulation, but the others were included as such high precipitation values are being observed increasingly in the UK (Cotterill et al., 2021). The storm precipitation depth was applied to each 25 x 25 m cell of Great Britain (GB). The model assumed all rain fell simultaneously and evenly across GB during a rain event, with instantaneous impacts. The depth of run-off accounting for sponginess (Q) was calculated at the 25 x 25 m resolution, and then arithmetically averaged per 1 km grid cell.

## 3. Application, Results and Discussion

### 3.1. Application

As far as we are aware, this study is the first to use the CN method to estimate water retention potential based on the sponginess of an entire country, using a point measure basis in a simple and effective way. Both the *I_a_* map and run-off maps are presented, as both are useful when assessing the ES value. The former indicates the sponginess of a grid square - i.e., the amount of precipitation in a single event that a grid square can ‘soak up’ based on the combination of geology and land management, while the latter shows the run-off after the application of a specific scenario. The results in the main text show the run-off given a 25 mm storm simulation, while Supplementary Information, S4 contains the comparative run-off results of the 50 mm and 75 mm storm simulations (Figs S4.1 and S4.2).

### 3.2. Results

The *I_a_* map (Fig. 2a) and the run-off map for the 25 mm storm simulation (Fig. 2b) indicate some distinctive patterns. The highest per-grid square run-off values were found in built-up, urban locations such as London and Birmingham, with almost 70% (17.25 / 25 mm) rainfall leaving those grid squares. Such city areas comprised mostly urban and sub-urban land covers, the former of which is highly impervious, with CNs between 89 – 95 and *I_a_* values of close to zero. With high levels of run-off from each grid square in cities, large volumes of water can accumulate at low lying points within them, leading to well documented instances of major flooding, especially in cities built around major rivers. (e.g., Jennings et al., 2015; Zhang et al., 2019; ARUP, 2023). Interestingly, some smaller towns had a similar level of run-off to cities, although the distribution of values was more varied within these towns, likely due to more urban / rural mixes of land cover types, and variation in slope. The Highlands region of Scotland generally exhibited the next highest levels of run-off, ranging between approximately 12.0 – 15.5 mm, which is between 48 and 62% of the precipitation delivered by a 25 mm storm event. This high run-off is primarily due to the extreme slopes of some of the mountains, such as Ben Nevis, which had a maximum slope of 193%. The effect of vegetation on run-off can be clearly distinguished in places like the Scottish Highlands, as individual grid squares that have the same slope can vary greatly in run-off depending on land cover type. For example, the mean run-off of bare rocks and coniferous woodland for a slope of 116% was 15.23 mm and 1.27 mm, respectively. Areas with the lowest run-off (<1 mm, 0 – 2%) had a very high proportion of the bog land cover type, including places such as the North Pennines, the North York Moors, and the Flow Country in Northern Scotland. In general, the most frequent run-off values were between 4 and 7 mm, indicating between 84% and 72% sponginess for a 25 mm rainfall event (Fig. 2). These areas generally coincided with *I_a_* values between 2 – 3 mm. See Supplementary Information, S5 (Figs S5.1 - S5.3), for extracted county-level images of Fig. 1, enabling differences to be more clearly observed.

**Fig. 1:**
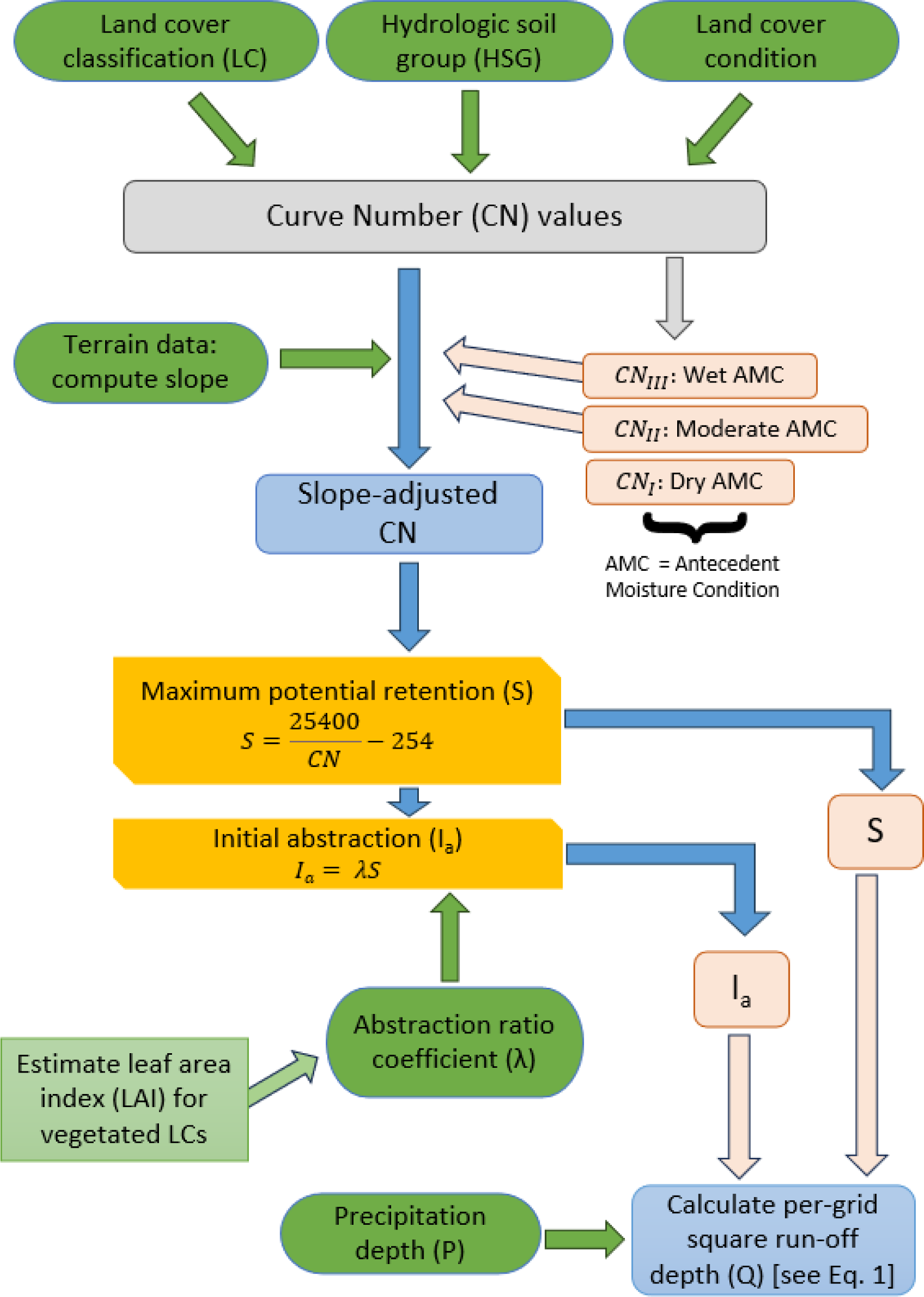
Schematic representation of the ‘sponginess’ model. The green shapes represent input requirements; yellow boxes represent the some of the processes of the model; brown-shapes represent the intermediate outputs of the model, which are subsequently fed back into the model; the green square box represents the choice we made for this study; and the blue boxes represents the outputs of the model.

**Fig. 2:**
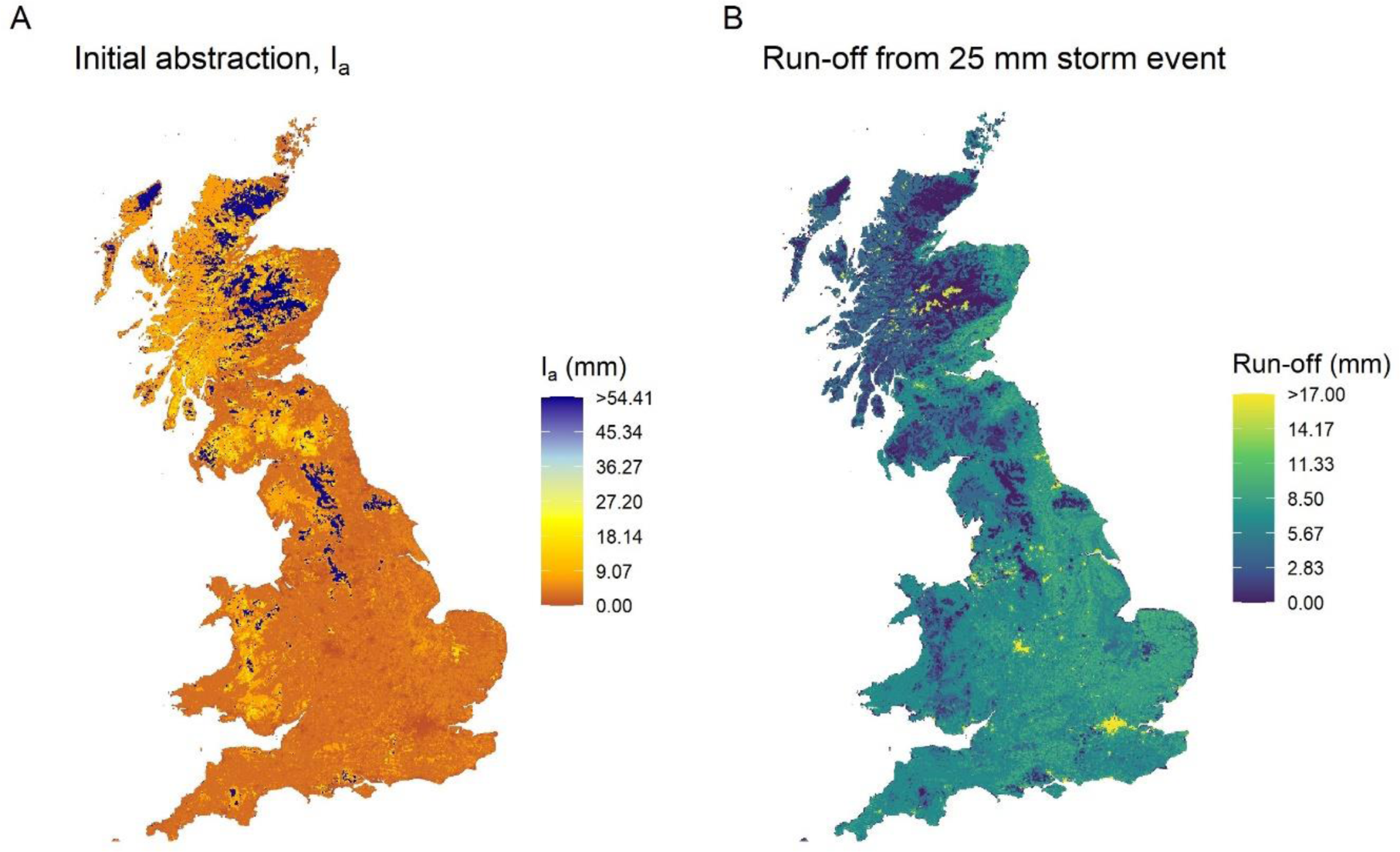
A) The average 1 km initial abstraction, *I_a_*, which is a function of maximum potential retention and rainfall interception proportion value; and B) average depth of estimated run-off across Great Britain at 1 km resolution given a 25 mm storm event. Both assumed a moderate antecedent moisture condition.

**Fig. 3:**
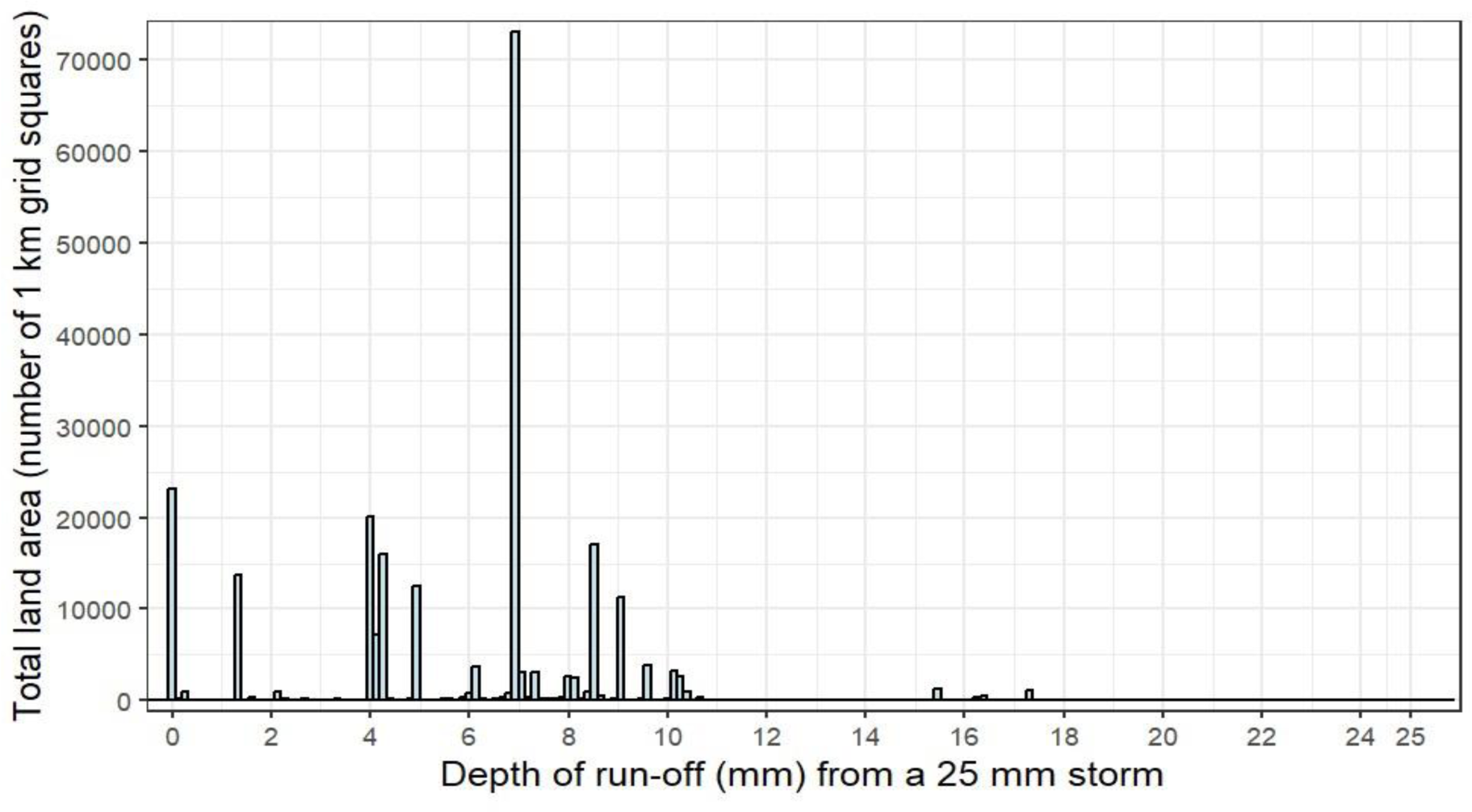
Histogram showing frequencies of the simulated run-off values per 1 km grid square after a 25 mm storm, over the whole of Great Britain.

### 3.3. Method applicability

It is important to note that the primary utility of this model lies in its ability to identify areas with a higher or lower potential for water retention and reduced run-off, rather than to estimate or calculate actual run-off or water retention. It can, therefore, be used to highlight vulnerabilities associated with land use change or opportunities for enhanced multifunctionality in a region. The model does not account for hydrological connectivity, such as upslope-downslope interactions, highlighting its role as a tool for identifying areas that contribute to high flood risk where further management may be necessary. While it does not have the functionality of comprehensive hydrological models, another potential use is that it could serve as a component within such models.

Modelling the results for sponginess on a grid basis (i.e., as a point measure) showed considerable variation across GB, enabling a high level of differentiation between neighbouring grid squares (Figs S5.1 – S5.3). Differentiation among grid squares was less in places with more homogenous, impervious land covers (e.g., cities). Combined, these outputs allow users to assess sponginess over large landscapes, but also to see the influence of an individual location on run-off. The underlying fine resolution of the land cover data, which was 25 x 25 m in this study, enabled more variation to be included when aggregated to the 1 km scale.

In terms of planning, having the results at a 1 km resolution – with each grid square treated as independent of its surroundings – would allow different management strategies to be simulated at a grid square level, and the results compared. For example, increasing the amount of ‘set-aside’ grassland can greatly alleviate flooding, but may come at the cost of reducing productive arable land, as documented by Evans and Boardman (2003). In the context of the CN method, the land use change described in Evans and Boardman (2003) would decrease the CN number by approximately half (63 to 30, assuming its HSG was A), increasing the sponginess. Crucially, the outcomes of any scenario can be compared with other simulated grid-based ES maps. This would aid in spatial prioritisation, providing stakeholders with a relative measure for policymaking and contributing to the development of ‘socially-relevant, process-based multifunctionality’ (Mastrangelo et al., 2014) of landscapes. It would also greatly aid trade-off analysis, especially in the context of planning scenarios, by allowing stakeholders to evaluate the relative benefits and costs of different land-use or management strategies on a range of ES, thereby making decisions that balance ecological and economic outcomes.

Moreover, and importantly, owing to the model’s simplicity, particularly its widely accessible inputs and ease-of-use, initial results and subsequent scenarios can be swiftly simulated. This would remove some of the issues that persist for organisations and nations that have limited access to resources or expertise, or a paucity of data (see e.g., Vigerstol and Aukema, 2011; Ruckelshaus et al., 2015; Willcock et al., 2016; Benra et al., 2021). However, there are limits to its applicability, therefore the decision on where and when to use the model should consider the limitations below.

### 3.4. User changes, limitations, and future improvements

The model presented here is adjustable in terms of its resolution of analysis, its extent, and its modelled time of year. Changing the extent and resolution would require adjustments to user-input elements that determine the original CN values, namely land cover classes, their condition, the soil type, and slope. If data are available for the scale of interest they can easily be used as the inputs, adding nuance to the results. Should data not be available or accessible at the desired resolution or extent, for example in data-scarce regions, then global, openly accessible data can be used. For instance, global data exist for land cover at a 100 m resolution (Buchhorn et al., 2020), 250 m resolution soil type (Ross et al., 2018; i.e., the HSG dataset used in this study), and high-resolution topographical elevation (e.g., Gesch et al., 1999). Assumptions would be required for land cover condition if these data were not available.

A key limitation of this study is the exclusion of dynamic weather conditions. This was a deliberate choice as the main purpose of this study was to demonstrate the utility of the slope-adjusted CN method in identifying how local land characteristics affect sponginess. This was best illustrated using a uniform precipitation scenario over the region, and as a snapshot in time, producing a spatially comparative measure. However, should one wish to model different times of year, incorporating variable weather conditions would be important. Dynamic weather conditions impact the distribution of precipitation both spatially and temporally. Moreover, precipitation events in the recent past influence antecedent moisture conditions (AMC) at a given location. Since both precipitation and AMC are critical inputs in determining landscape sponginess, their accurate representation is essential for temporally robust model outcomes. Due to the land’s sponginess, input precipitation data that are higher or lower than the data used here will affect the final run-off in a corresponding, but non-linear, manner, compared to the modelled results (Fig 2), an example of which can be seen in Supplementary Information, S4. However, it is important to note greater uncertainty can be expected in CN model results when precipitation levels are increased (Deshpande and Amit Dhorde, 2024). Additionally, how much interception trees provide during heavy rain events is still debated (Page et al., 2020; Cooper et al., 2021).

The other major input affected by time of year is the combination of land cover and *λ*, an adjustable constituent of *I_a_*. Here, to parameterise *λ*, due to the national-scale approach, we primarily relied on LAI, which is a well-known vegetated land use parameter. Since most LAI studies focus on fully-grown or mature crops, we approximated winter LAIs. However, LAI, particularly in arable land, varies greatly through and between seasons, as well as due to management practices (e.g., Dammer et al., 2008; Sieling et al., 2016). Consequently, ‘sponginess’ maps would differ greatly depending on the time of year. For instance, we reduced winter crops’ LAI by 75% from their literature-derived average LAI, thereby reducing the initial abstraction of areas with those crops. Future adjustments to input LAI could account for expected crop growth stage corresponding to the modelled time of year. Alternatively, *λ* could be simplified by using a single region-specific value regardless of land cover type, which is a common method for parameterising *λ* (Ling and Yusop, 2014). This approach would still yield relative values for water retention potential, as sponginess would continue to be influenced by other landscape factors. This list is non-exhaustive as a way of parametrising *I_a_*, as much debate exists around its parametrisation (Jain et al., 2006; Wang et al., 2012; Ajmal et al., 2016).

An omission from this model is human interventions, such as engineered flood management infrastructure, which could influence run-off. For example, the uptake of rain gardens in cities limits the impacts of stormwater, and can have significant effects on localised sponginess (Jennings et al., 2015). Model validation has also not been conducted due to challenges in validating independent grid square estimates, which only represent part of the hydrological system. Previous validation of the CN methodology has involved small-scale comparisons between observed run-off and CN estimations (e.g., Deshpande and Amit Dhorde, 2024) or hydrologic models like SWAT (Arnold et al., 2012) and HEC-HMS (Feldman, 2000), which consider hydrological connectivity, not point estimates. Nonetheless, realistic relative variations across regions can be seen in Figs S5.1 – S5.3, verifying expected trends. These are both important considerations, particularly when modelling landscape with a high urban and sub-urban composition where the default CN values are high.

### 3.5. Conclusion

This research introduces a model to estimate water retention, a key regulating ecosystem service, at a management relevant resolution (parcel scale). By applying it to Great Britain we demonstrate how the model can help identify locations where additional management could reduce flood risk or where further enquiry using more complex hydrological models might be beneficial. The model is versatile and its simplicity ensures that it requires minimal resources and expertise, making it accessible in a variety of situations.

## 4. Code availability

Code to run the model are available on request from the corresponding author (PME).

## 5. CRediT authorship contribution statement

**Paul M. Evans**: Conceptualization, Data curation, Formal analysis, Methodology, Writing – original draft. **Alejandro Dussaillant**: Conceptualization, Methodology, Writing – review & editing. **Varun Varma**: Conceptualization, Methodology, Writing – review & editing. **John W. Redhead**: Conceptualization, Writing – review & editing. **Richard F. Pywell**: Conceptualization, Funding acquisition, Project administration, Writing – review & editing. **Jonathan Storkey**: Conceptualization, Funding acquisition, Project administration, Writing – review & editing. **Andrew Mead**: Conceptualization, Methodology, Writing – review & editing **James M. Bullock**: Conceptualization, Methodology, Writing – review & editing, Supervision.

## 6. Declaration of competing interest

The authors declare that they have no known competing financial interests or personal relationships that could have appeared to influence the work reported in this paper.

## 7. Acknowledgement

We would like to thank the AgLand project team for co-ordination help and constructive discussion and advice. This work was funded by UKRI Grants NE/T000244/2, NE/T001178/1, and 10093115.

## Notes

### Competing Interest Statement

The authors have declared no competing interest.

