## Supplementary material for "Quantifying relative sponginess: a high-resolution model of landscape water retention as an ecosystem service": Sponginess_SI_final_20240930.docx

Contents:

S1: Antecedent moisture condition (AMC)

S2: Adjustment values for factors that affect roughness of flood plains.

S3: Leaf area index trends for each vegetation type

S4: Run-off from different storms

S5: Sponginess in different regions

Supplementary Information references

#### S1: Antecedent moisture condition (AMC)

Ponce and Hawkins (1996) noted, building upon work of USDA (1986) and Hawkins et al. (1985), how there can be variability around the mean CN, and that this can be considered in the calculations. Specifically, they used a metric indicating the inherent fluctuations in soil moisture and its correlation with rainfall-runoff dynamics: AMC. The standard AMC was, and is generally (as in this study), used to derive *CN_II_*, which assumes that the prior soil conditions before a rainfall event were not saturated or arid. However, for values of CN between 55 and 95, two other values can be used, indicating different prior soil conditions. CN*_I_*, representing the lowest run-off potential due to prior dry conditions (eq. A1), and CN*_III_*, representing the highest potential run-off due to prior wet conditions (eq. A2) were calculated using the equations of Sharpley and Williams (1990):

$$\left( eq.A1 \right){CN}_{I}=\frac{{CN}_{II}}{2.281-0.01281{CN}_{II}}$$

$$\left( eq.A2 \right){CN}_{III}=\frac{{CN}_{II}}{0.427+0.00573{CN}_{II}}$$

where *CN_II_* is the average run-off potential (i.e., the one applied with medium prior wetness conditions). This can result in the CN number being different by up to 20 (Table A1).

Table S1.1: Comparisons between CN based on the prior AMC condition.

| CN*_II_* | CN*_I_* | CN*_III_* |
| --- | --- | --- |
| 55 | 34.88852 | 74.10901 |
| 60 | 39.67204 | 77.8412 |
| 65 | 44.87866 | 81.3059 |
| 70 | 50.56707 | 84.53085 |
| 75 | 56.80742 | 87.54012 |
| 80 | 63.68413 | 90.35464 |
| 85 | 71.29975 | 92.99272 |
| 90 | 79.78016 | 95.47046 |
| 95 | 89.28152 | 97.80203 |

#### S2: Adjustment values for factors that affect roughness of flood plains.

Table S2.1: Adjustment values for factors that affect roughness of flood plains. Reproduced from Acrement and Schneider (1984).

| Flood plain vegetation | Roughness adjustment value | Example |
| --- | --- | --- |
| Small | 0.001-0.010 | Dense growth of flexible turf grass, such as Bermuda, or weeds growing where the average depth of flow is at least two times the height of the vegetation, or supple tree seedlings such as willow, cottonwood, arrowweed, or saltcedar growing where the average depth of flow is at least three times the height of the vegetation. |
| Medium | 0.010-0.025 | Turf grass growing where the average depth of flow is from one to two times the height of the vegetation, or moderately dense stemmy grass, weeds, or tree seedlings growing where the average depth of flow is from two to three times the height of the vegetation; brushy, moderately dense vegetation, similar to 1- to 2-year-old willow trees in the dormant season. |
| Large | 0.025-0.050 | Turf grass growing where the average depth of flow is about equal to the height of the vegetation, or 8- to 10-year-old willow or cottonwood trees intergrown with some weeds and brush (none of the vegetation in foliage) where the hydraulic radius exceeds [0.61 m], or mature row crops such as small vegetables, or mature field crops where depth of flow is at least twice the height of the vegetation. |
| Very large | 0.050-0.100 | Turf grass growing where the average depth of flow is less than half the height of the vegetation, or moderate to dense brush, or heavy stand of timber with few down trees and little undergrowth where depth of flow is below branches, or mature field crops where depth of flow is less than the height of the vegetation. |
| Extreme | 0.100-0.200 | Dense bushy willow, mesquite, and saltcedar (all vegetation in full foliage), or heavy stand of timber, few down trees, depth of flow reaching branches. |

#### S3: Leaf area index trends for each vegetation type

Table S3.1: Leaf area index (LAI) for average European land cover types.

| Land cover | Acrement and Schneider value (max = 0.2) | Original LAI | Reference | Modifying effect | Winter proxy LAI | Calculated λ |
| --- | --- | --- | --- | --- | --- | --- |
| Broadleaf woodland | 0.1 | 4.74 | Munier et al., 2018 | 50% of original LAI | 2.37 | 0.12 |
| Coniferous woodland | 0.2 | 4.65 | Munier et al., 2018 | No | 4.65 | 0.26 |
| Winter wheat [winter sown] | 0.025 - 0.1 | 3.50 | Gebbers et al., 2011 | 75% of original LAI | 2.63 | 0.14 |
| Winter barley [winter sown] |  | 2.38 | Rosso et al., 2022 | 75% of original LAI | 1.79 | 0.08 |
| Oil seed rape [winter sown] |  | 3.38 | Zhang et al., 2023; Ploschuk et al., 2021 | 75% of original LAI | 2.53 | 0.13 |
| Spring wheat [spring sown] |  | 2.18 | Dong et al., 2019 | 50% of original LAI | 1.09 | 0.04 |
| Oat  [spring sown] |  | 2.50 | Peltonen-Sainio et al., 1997 | 50% of original LAI | 1.25 | 0.05 |
| Maize [spring sown] |  | 3.40 | González-Sanpedro et al., 2008 | 25% of original LAI | 0.85 | 0.03 |
| Spring barley [spring sown] |  | 1.84 | Rosso et al., 2022 | 50% of original LAI | 0.92 | 0.03 |
| Potato [spring sown] |  | 5.40 | González-Sanpedro et al., 2008 | 25% of original LAI | 1.35 | 0.06 |
| Beans [spring sown] |  | 4.2 | Setiyono et al., 2008 | 50% of original LAI | 2.10 | 0.10 |
| Other arable |  | 3.20 | Average of all crop categories | 50% of original LAI | 1.60 | 0.07 |
| Arable grass |  | 2.54 | Dusseux et al., 2014; Kang et al., 2016 | 50% of original LAI | 1.27 | 0.05 |
| Improved grassland | 0.01 | 2.54 | Dusseux et al., 2014; Kang et al., 2016 | 50% of original LAI | 1.27 | 0.05 |
| Neutral grassland |  | 3.26 | Klingler et al., 2020; Darvishzadeh et al., 2008; Atzberger et al., 2015; Munier et al., 2018 | 75% of original LAI | 2.44 | 0.12 |
| Calcareous grassland |  | 3.26 | Klingler et al., 2020; Darvishzadeh et al., 2008; Atzberger et al., 2015; Munier et al., 2018 | 75% of original LAI | 2.44 | 0.12 |
| Acid grassland |  | 3.26 | Klingler et al., 2020; Darvishzadeh et al., 2008; Atzberger et al., 2015; Munier et al., 2018 | 75% of original LAI | 2.44 | 0.12 |
| Heather grassland |  | 3.26 | Klingler et al., 2020; Darvishzadeh et al., 2008; Atzberger et al., 2015; Munier et al., 2018 | 75% of original LAI | 2.44 | 0.12 |
| Heather | 0.011-0.025 | 3.26 | Value taken from ‘heather grassland’ |  | 2.44 | 0.12 |
| Fen, marsh and swamp | N/A | N/A | N/A | No | N/A | 0.95 |
| Bog |  |  |  |  |  | 0.95 |
| Inland rock |  |  |  |  |  | 0.08 |
| Saltwater |  |  |  |  |  | 0.95 |
| Freshwater |  |  |  |  |  | 0.95 |
| Supralittoral rock |  |  |  |  |  | 0.08 |
| Supralittoral sediment |  |  |  |  |  | 0.4 |
| Littoral rock |  |  |  |  |  | 0.08 |
| Littoral sediment |  |  |  |  |  | 0.4 |
| Saltmarsh |  |  |  |  |  | 0.95 |
| Urban |  |  |  |  |  | 0.01 |
| Suburban |  |  |  |  |  | 0.03 |

#### S4: Run-off from different storms

This section includes the results for the simulation described in the main text. An important thing to note is that as the event depth increases, there would likely be less retention and higher run-off from areas upslope of each grid square (which were not considered in this model), so the uncertainty of predictions is higher for larger rainfall events (e.g., Deshpande and Amit Dhorde., 2024).

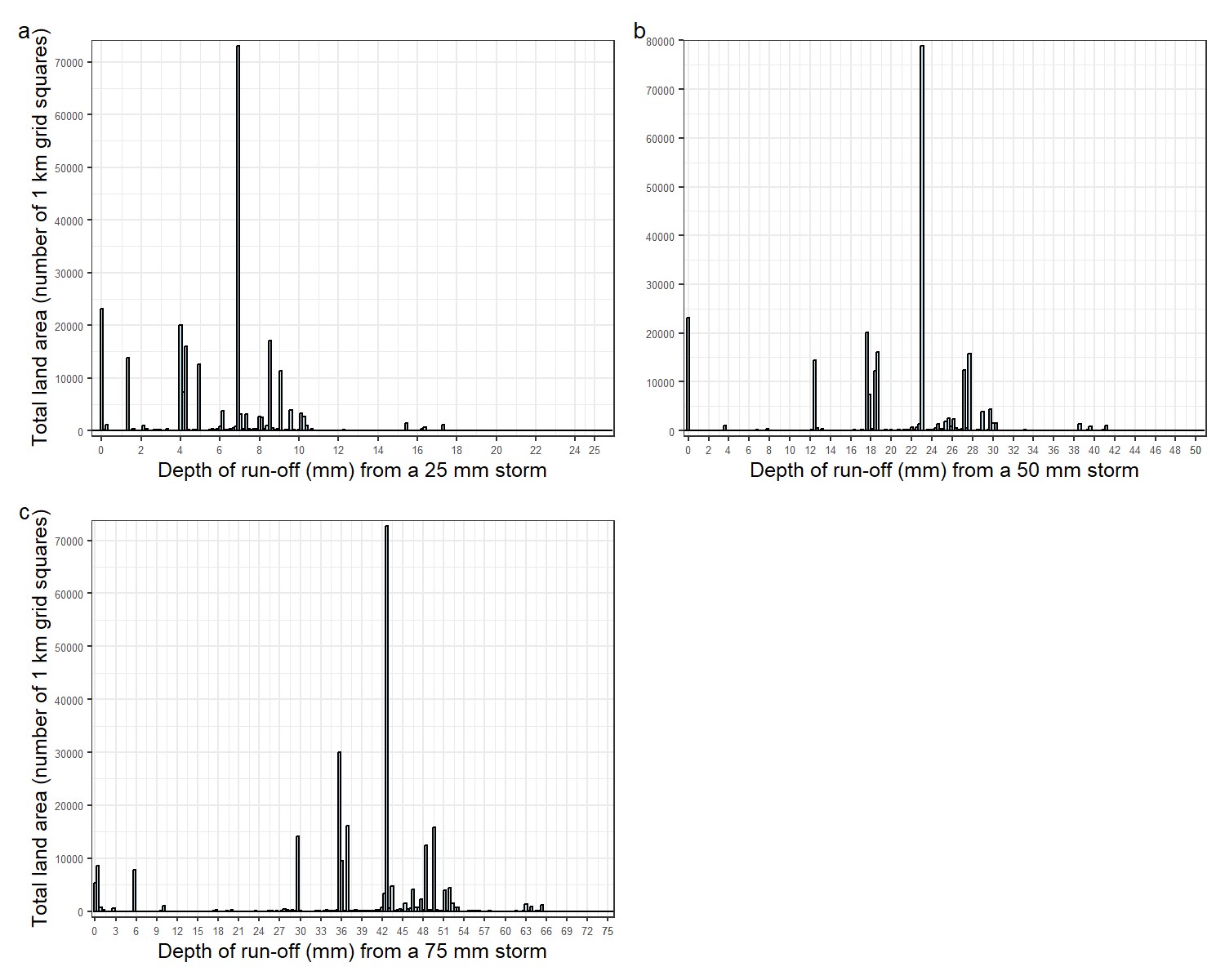
Fig S4.1: a) Depth of run-off across Great Britain at 1 km resolution after a a) 25 mm storm event; b) 50 mm storm event; and c) 75 mm storm event, over the whole of Great Britain. The graphs show the depth of run-off.

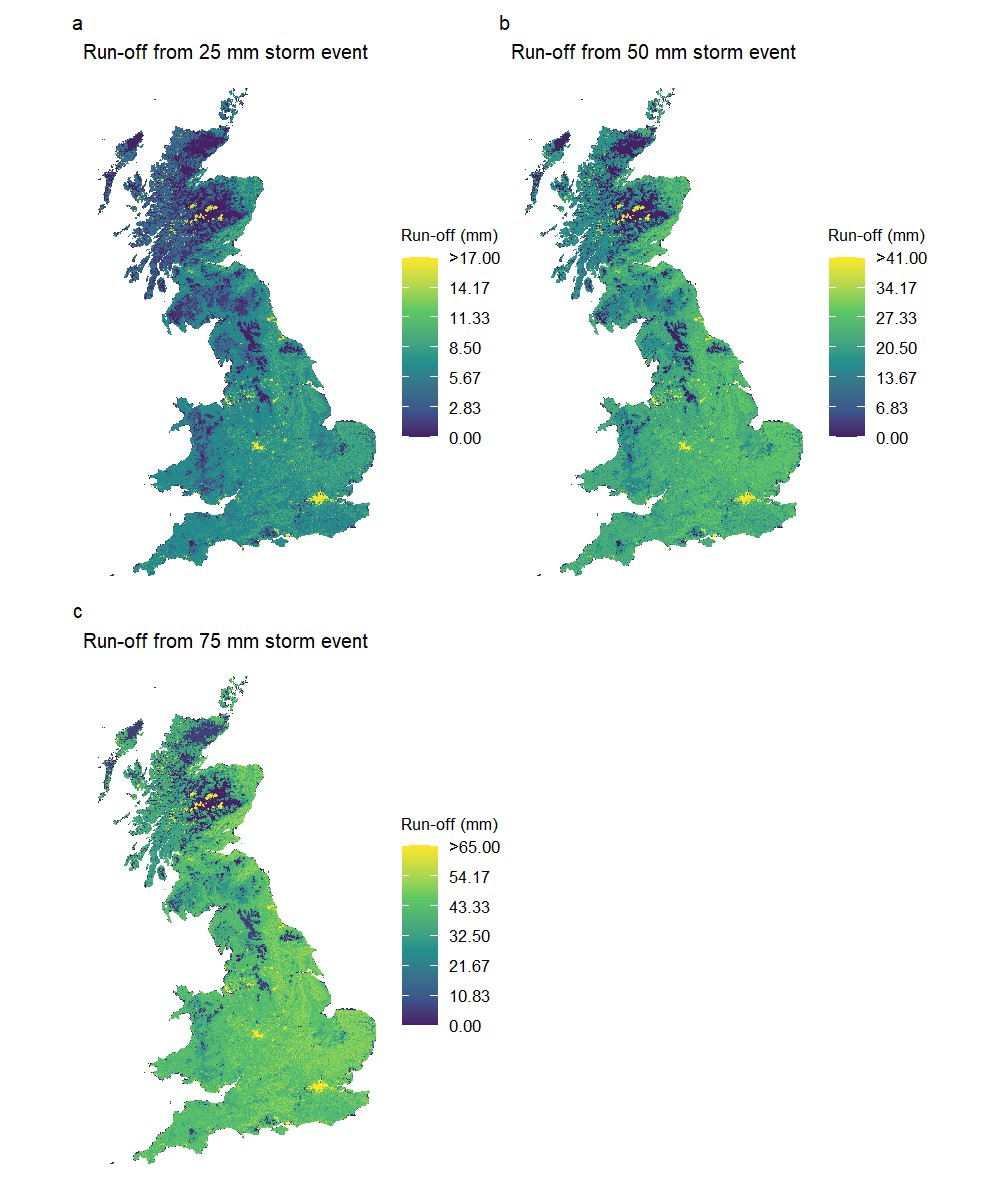

Fig S4.2: Depth of run-off across Great Britain at 1 km resolution after a a) 25 mm storm event; b) 50 mm storm event; and c) 75 mm storm event.

#### S5: Sponginess in different regions

This section shows the results from Figs 2a and 2b but only for select regions of the UK, namely the counties of Greater London, Inverness, and Mid Glamorgan.

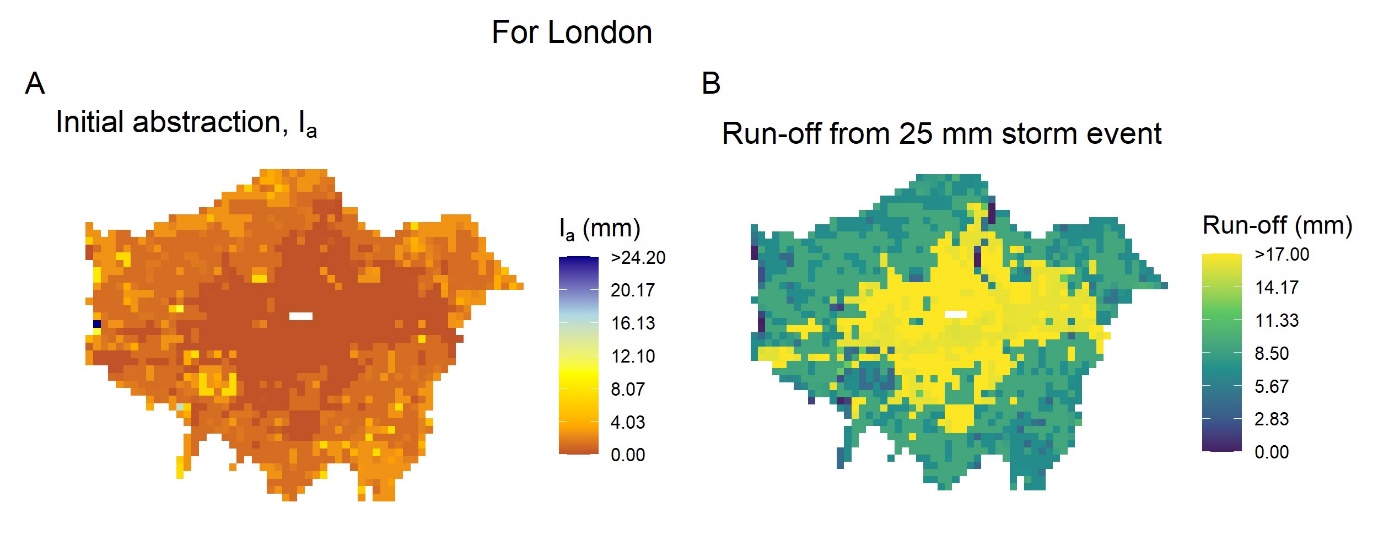
Fig S5.1: Sponginess maps for London. A) The average 1 km initial abstraction, *I_a_*, which is a function of maximum potential retention and rainfall interception proportion value; and B) average depth of estimated run-off across Great Britain at 1 km resolution given a 25 mm storm event. Both assumed a moderate antecedent moisture condition.

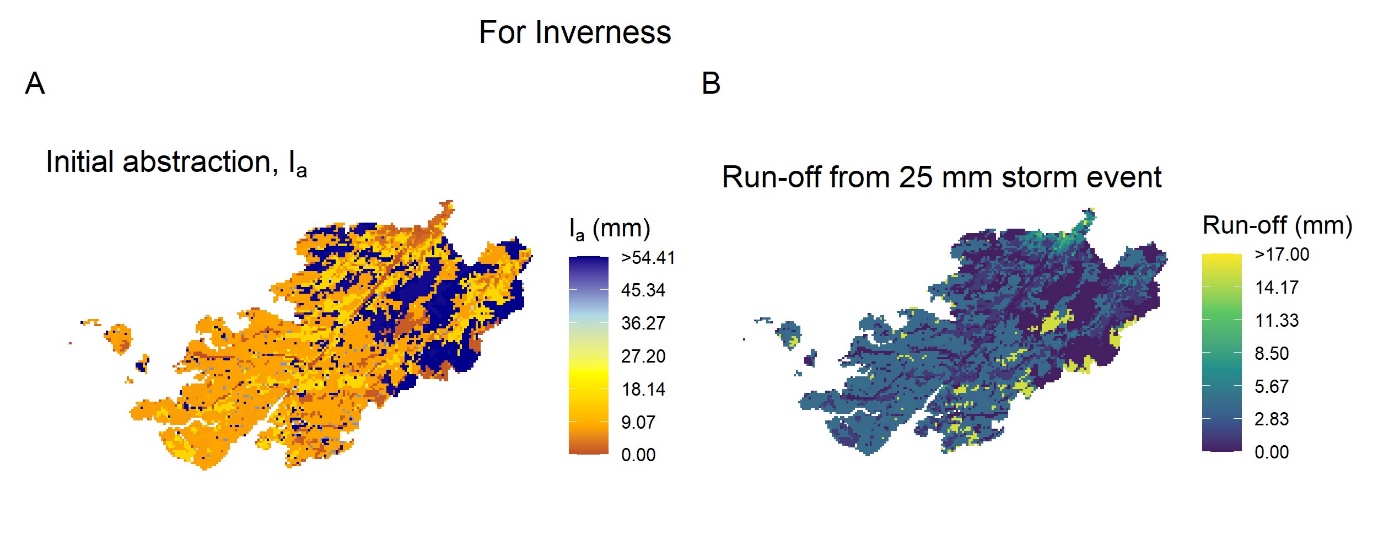
Fig S5.2: Sponginess maps for Inverness. See full caption to Fig A3 for details.

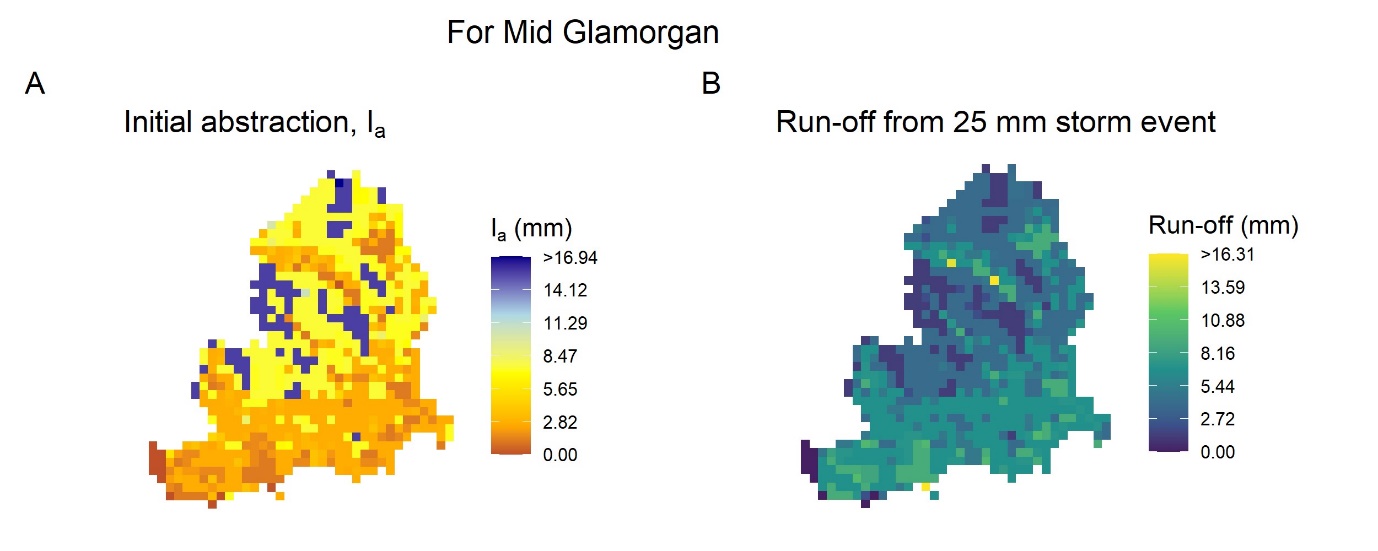
Fig S5.3: Sponginess maps for Mid Glamorgan. See full caption to Fig A3 for details.

### Supplementary Information references

Acrement, G.J. & Schneider, V.R. (1984). *Guide for selecting Manning’s roughness coefficients for natural channels and floodplains* (FHWA-TS-84-204). Federal Highways Administration US Department of Transportation. <https://pubs.usgs.gov/wsp/2339/report.pdf>

Atzberger, C., Darvishzadeh, R., Immitzer, M., Schlerf, M., Skidmore, A., & le Maire, G. (2015). Comparative analysis of different retrieval methods for mapping grassland leaf area index using airborne imaging spectroscopy. *International Journal of Applied Earth Observation and Geoinformation*, *43*, 19–31. <https://doi.org/10.1016/j.jag.2015.01.009>

Darvishzadeh, R., Skidmore, A., Schlerf, M., Atzberger, C., Corsi, F., & Cho, M. (2008). LAI and chlorophyll estimation for a heterogeneous grassland using hyperspectral measurements. *ISPRS Journal of Photogrammetry and Remote Sensing*, *63*(4), 409–426. <https://doi.org/10.1016/j.isprsjprs.2008.01.001>

Deshpande, G., & Amit Dhorde, A. (2024). Validating the Curve Number estimation approaches: A case study of an urbanizing watershed from Western Maharashtra, India. *Modeling Earth Systems and Environment*, *10*(2), 1615–1629. <https://doi.org/10.1007/s40808-023-01855-7>

Dong, T., Liu, J., Shang, J., Qian, B., Ma, B., Kovacs, J. M., Walters, D., Jiao, X., Geng, X., & Shi, Y. (2019). Assessment of red-edge vegetation indices for crop leaf area index estimation. *Remote Sensing of Environment*, *222*, 133–143. <https://doi.org/10.1016/j.rse.2018.12.032>

Dusseux, P., Gong, X., Hubert-Moy, L., & Corpetti, T. (2014). Identification of grassland management practices from leaf area index time series. *Journal of Applied Remote Sensing*, *8*(1), 083559. <https://doi.org/10.1117/1.JRS.8.083559>

Gebbers, R., Ehlert, D., & Adamek, R. (2011). Rapid Mapping of the Leaf Area Index in Agricultural Crops. *Agronomy Journal*, *103*(5), 1532–1541. <https://doi.org/10.2134/agronj2011.0201>

González-Sanpedro, M. C., Le Toan, T., Moreno, J., Kergoat, L., & Rubio, E. (2008). Seasonal variations of leaf area index of agricultural fields retrieved from Landsat data. *Remote Sensing of Environment*, *112*(3), 810–824. <https://doi.org/10.1016/j.rse.2007.06.018>

Hawkins, R., Hjelmfelt, A. T., & Zevenbergen, A. W. (1985). Runoff Probability, Storm Depth, and Curve Numbers. *Journal of Irrigation and Drainage Engineering*, *111*(4), 330–340. <https://doi.org/10.1061/(ASCE)0733-9437(1985)111:4(330)>

Kang, Y., Özdoğan, M., Zipper, S. C., Román, M. O., Walker, J., Hong, S. Y., Marshall, M., Magliulo, V., Moreno, J., Alonso, L., Miyata, A., Kimball, B., & Loheide, S. P. (2016). How Universal Is the Relationship between Remotely Sensed Vegetation Indices and Crop Leaf Area Index? A Global Assessment. *Remote Sensing*, *8*(7), Article 7. <https://doi.org/10.3390/rs8070597>

Klingler, A., Schaumberger, A., Vuolo, F., Kalmár, L. B., & Pötsch, E. M. (2020). Comparison of Direct and Indirect Determination of Leaf Area Index in Permanent Grassland. *PFG – Journal of Photogrammetry, Remote Sensing and Geoinformation Science*, *88*(5), 369–378. <https://doi.org/10.1007/s41064-020-00119-8>

Munier, S., Carrer, D., Planque, C., Camacho, F., Albergel, C., & Calvet, J.-C. (2018). Satellite Leaf Area Index: Global Scale Analysis of the Tendencies Per Vegetation Type Over the Last 17 Years. *Remote Sensing*, *10*(3), Article 3. <https://doi.org/10.3390/rs10030424>

Peltonen-Sainio, P., Forsman, K., & Poutala, T. (1997). Crop Management Effects on Pre-and Post-Anthesis Changes in Leaf Area Index and Leaf Area Duration and their Contribution to Grain Yield and Yield Components in Spring Cereals. *Journal of Agronomy and Crop Science*, *179*(1), 47–61. <https://doi.org/10.1111/j.1439-037X.1997.tb01146.x>

Ploschuk, R. A., Miralles, D. J., & Striker, G. G. (2021). Early- And late-waterlogging differentially affect the yield of wheat, barley, oilseed rape and field pea through changes in leaf area index, radiation interception and radiation use efficiency. *Journal of Agronomy and Crop Science*, *207*(3), 504–520. <https://doi.org/10.1111/jac.12486>

Ponce, V. M., & Hawkins, R. H. (1996). Runoff Curve Number: Has It Reached Maturity? *Journal of Hydrologic Engineering*, *1*(1), 11–19. <https://doi.org/10.1061/(ASCE)1084-0699(1996)1:1(11)>

Rosso, P., Nendel, C., Gilardi, N., Udroiu, C., & Chlebowski, F. (2022). Processing of remote sensing information to retrieve leaf area index in barley: a comparison of methods. *Precision Agriculture*, *23*(4), 1449–1472. <https://doi.org/10.1007/s11119-022-09893-4>

Setiyono, T. D., Weiss, A., Specht, J. E., Cassman, K. G., & Dobermann, A. (2008). Leaf area index simulation in soybean grown under near-optimal conditions. *Field Crops Research*, *108*(1), 82–92. <https://doi.org/10.1016/j.fcr.2008.03.005>

USDA. (1986). *Urban Hydrology for Small Watersheds* (TR-55). Natural Resources Conservation Service.

Zhang, W., Li, Z., Pu, Y., Zhang, Y., Tang, Z., Fu, J., Xu, W., Xiang, Y., & Zhang, F. (2023). Estimation of the Leaf Area Index of Winter Rapeseed Based on Hyperspectral and Machine Learning. *Sustainability*, *15*(17), Article 17. <https://doi.org/10.3390/su151712930>
